# Bacteriophage genomics: What has five years of INPHARED taught us?

**DOI:** 10.64898/2026.05.06.722914

**Authors:** Ryan Cook, Branko Rihtman, Alise J. Ponsero, Slawomir Michniewski, Andrea Telatin, Thomas Sicheritz-Pontén, Evelien M. Adriaenssens, Andrew D. Millard

## Abstract

Bacteriophages are key drivers of microbial ecology and evolution, and the rapid expansion of phage sequencing has created sustained demand for curated reference genome databases. We released the INfrastructure for a PHAge REference Database (INPHARED) in January 2021 to provide quality-controlled metadata for complete phage genomes from cultured isolates. Here, we compare the 2021 and 2026 snapshots, spanning a five-year period that included a substantial overhaul of bacterial virus taxonomy by the ICTV. The database has approximately doubled, from 14,244 to 28,777 genomes, yet the proportion representing novel species-level diversity has declined, indicating that redundant sequencing is outpacing new discovery. Host bias persists despite the addition of 97 new host genera. We have incorporated genome quality assessments, lifestyle predictions, and defence and anti-defence system annotations, providing an updated resource and a snapshot of the current state of phage genomics.

## Background

In all environments where they’ve been studied in detail, bacteriophages are known to be of significant ecological consequence. Bacteriophages, or simply phages, are the obligate intracellular parasites of bacteria. Phages drive bacterial evolution, both as microbial predators and agents of horizontal gene transfer^1^. Our understanding of phages, and their diverse virus-host interactions, has massively expanded since the sequencing of φX174 in 1977^1^.

Now, approaching 50 years later, the ease of high-throughput sequencing and renewed interest in phage therapy—largely due to the emergence of antimicrobial resistance—has led to ever increasing numbers of phage genomes being sequenced ^2,3^. In turn, this has led to increased demand and need for curated reference databases, providing quality-controlled metadata and a standardised framework against which new genomes can be compared.

In January 2021, we released the INfrastructure for a PHAge REference Database (INPHARED)^4^, combining automated retrieval, quality control and standardised annotation of complete phage genomes. The database was focused on complete phage genomes from cultured isolates, excluding prophage predictions and the vast majority of metagenomically derived sequences. Curated reference databases are foundational to phage genomics research, providing quality-controlled metadata and a standardised framework against which new genomes can be compared. The data provided by INPHARED has been incorporated into a number of tools used for the analysis of phage genomes, including Pharokka^4^, Phold^5^, and *taxMyPhage*^6^, and have served as reference datasets for vConTACT2 analyses^7^ and MASH-based genome comparisons^7^.

Furthermore, curated datasets such as INPHARED are essential for benchmarking in the development of machine learning approaches for phage-host interaction prediction^8,9^ and taxonomic assignments^10^. This breadth of use underscores the importance of maintaining a well-curated, regularly updated reference database for the phage genomics community.

In the five years since the original publication, several developments in the field have necessitated updates to the INPHARED database and prompted a re-examination of the state of phage sequencing. The International Committee on Taxonomy of Viruses (ICTV) Bacterial Viruses Subcommittee introduced substantial changes to the taxonomy of bacterial viruses. The morphology-based families *Myoviridae*, *Siphoviridae*, and *Podoviridae*, along with the order *Caudovirales*, were abolished in favour of genomically defined taxa^11,12^. This change was accompanied by a dramatic expansion in the number of recognised taxa (orders, families, genera, and species), reflecting the surge in sequenced phage genomes from isolates and metagenomes. In parallel, there has been an explosion of interest in the “arms race” between bacterial defence systems and phage counter-defence strategies^13–16^, with tools such as DefenseFinder and PADLOC now enabling the systematic annotation of these systems across large bacterial genome collections^17,18^.

Methodological advances in phage isolation and sequencing technologies also continue to shape the field. The use of robotics and miniaturisations of phage isolation techniques is increasing the throughput of phage that are been isolated^19^, along with innovative approaches such as Phage DisCo, which has been used to massively expand the number of isolated *Tectiviridae*^20,21^. Sequencing technologies have further developed with increased accuracy from ONT sequencing and increased throughput from PacBio sequencing. These long-read technologies offer both the ability to recover increasing number of complete phage genomes from metagenomes and the ability to identify modified bases in phage genomes^22–27^. The portability of ONT minION allows phages to be sequenced by research labs rather than commercial suppliers, increasing the number of phage genomes that can be sequenced. ONT sequencing has also facilitated recent developments allowing individual plaques to be sequenced rather than large scale lysates, which will likely further increase the number of phage genomes sequenced^28,29^.

With ever increasing numbers of phage genomes and the aforementioned changes to the field, we have redeveloped the INPHARED codebase to allow faster updates to the database and incorporated further metadata, prediction of defence systems, more comprehensive taxonomy and lifestyle predictions. Here, we compare the 2021 and 2026 datasets to assess how the landscape of phage genomics has changed over this period, including newly integrated data on updated taxonomy, defence and anti-defence system annotations, genome completeness assessments, and lifestyle predictions.

## Methods

The original code for INPHARED has now been rewritten in python and is available at XXX. The same selection criteria for identification of phage genomes have been maintained. Taxonomy of genomes was predicted by use of taxMyPhage v0.3.4^30^, using the VMRv40. Phage lifestyle was predicted using PhaTYP as part of PhaBOX v2.1.10^31^. Genome quality assessment was performed using CheckV v1.0.3^32^. Phage-encoded defence and anti-defence systems were identified using DefenseFinder v2.0.1^17^.

Genome clustering was performed using BLAST v2.16.0+ with the anicalc.py and aniclust.py scripts available with CheckV (https://bitbucket.org/berkeleylab/checkv/src)^32–34^. Antimicrobial resistance genes and virulence factors were predicted using ABRicate v1.2.0 (https://github.com/tseemann/ABRICATE) with NCBI AMRFinderPlus^35^ and VFDB^36^ databases. All plots were produced in R Studio v4.2.2^37^ using ggplot2^38^.

## Results and Discussion

### The database has doubled, but redundancy is increasing

As of February 2026, INPHARED contains 28,777 complete phage genomes, up from 14,244 in January 2021, approximately a doubling over five years (Figure 1A). Of the original 14,244 accessions, 13,738 are retained in the 2026 dataset and 506 have been removed, with 15,039 new accessions added. Most removals (348) were genomes that did not meet our inclusion criteria of complete genomes of phage isolates, such as satellite phages, predominantly associated with *Streptococcus*, which were excluded due to lacking autonomous virion structural genes ^39^, with the remainder reflecting community curation.

**Figure 1.**
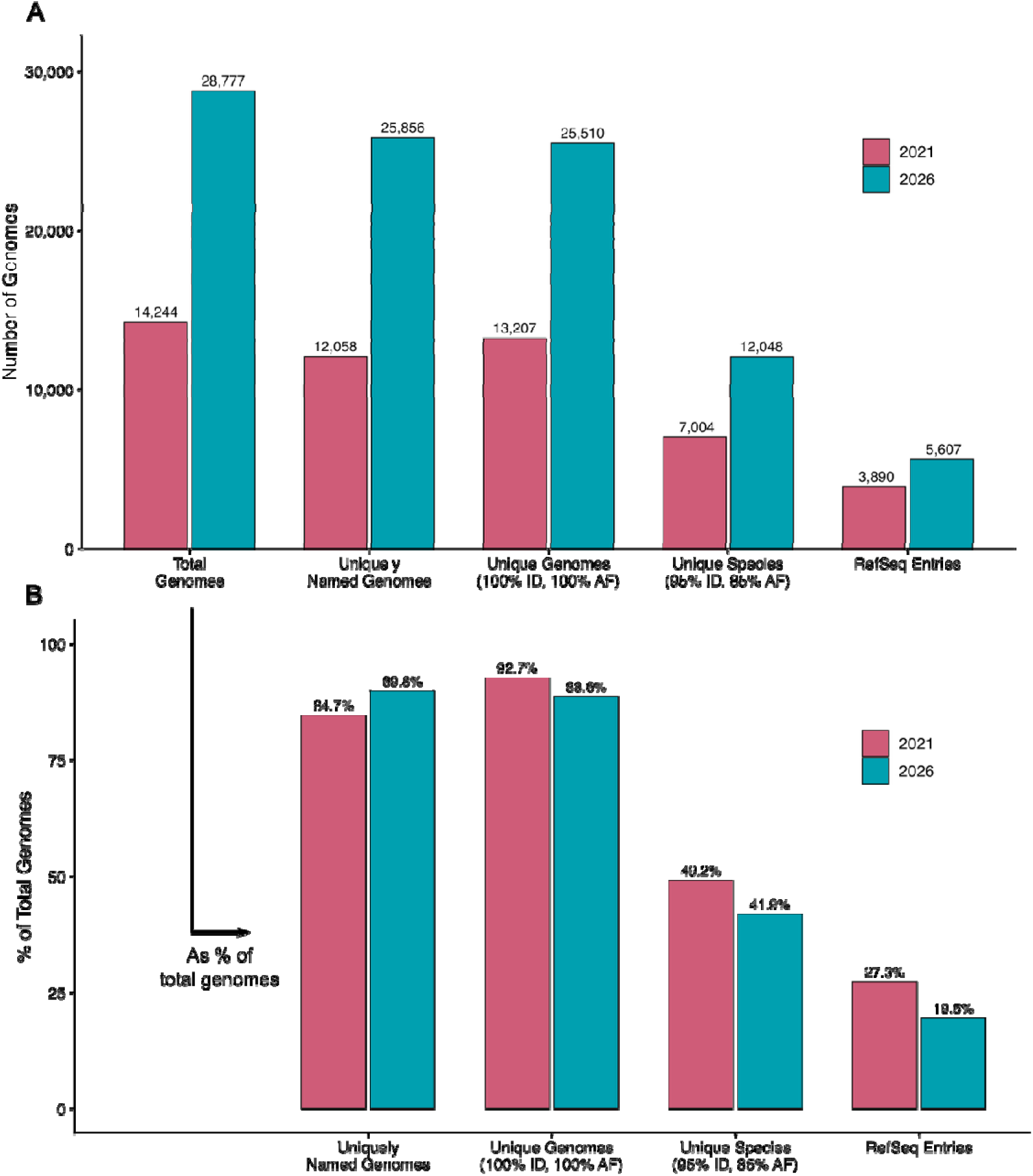
Database Increase in 5 Years. Paired bar plots comparing the 2021 (left) and 2026 (right) datasets, with number of genomes (**A**) and proportion of total genomes (**B**).

However, these 28,777 accessions do not all represent unique phages. Only 25,856 accessions have unique GenBank names, with many duplications and inconsistencies or uninformative names (e.g. “Microviridae sp.” [n=998], “Enterobacteria phage ID11” [n=339], and “Bacteriophage sp.” [n=51x]). When de-replicated at 100% average nucleotide identity (ANI) over 100% alignment of the shorter sequence, the dataset reduced to 25,510 unique genomes. Applying MIUViG species-level thresholds (95% ANI over 85% alignment of the shorter sequence^40^), resulted in 12,048 viral operational taxonomic units (vOTUs). Of the 28,777 accessions, only 5,607 have a corresponding RefSeq record. These values have approximately doubled since 2021, when INPHARED contained 13,207 unique genomes, 7,004 vOTUs, and 3,890 RefSeq records from 14,244 total accessions.

Whilst the absolute numbers have grown, the proportions tell a more nuanced story. The proportion of accessions representing a unique genome has decreased from 92.7% to 88.6%, and the proportion representing a unique species-level vOTU has fallen from 49.2% to 41.9%, indicating that the rate of redundant sequencing is outpacing novel phage discovery (Figure 1B). The proportion of accessions with a corresponding RefSeq record has also decreased, from 27.3% to 19.5%. Conversely, the proportion of accessions with a unique name has increased from 84.7% to 89.8%, suggesting an improvement in naming practices^41^.

### Isolation host metadata is inconsistently reported

The 2026 dataset introduces a new “Isolation Host” field, extracted from GenBank records for all 28,777 genomes (5,418 unique values), often providing strain-level resolution and insight into host bias. For example, *Mycobacterium smegmatis* mc²155 is recorded as the isolation host for 2,347 genomes but appears under 32 inconsistent entries due to typographical errors (e.g. “Mycobacteruim”), formatting differences (e.g. “mc2 155”, “mc#155”, “mc2155”, “mc(2)155”), and reclassification (e.g. *Mycolicibacterium)*. This highlights a lack of standardisation in recording host information and raises challenges around handling bacterial taxonomy. Within the INPHARED metadata we have automated the correction of the most common typographical errors for bacterial genera and merging of obvious variants of the same strain name into a single common name. Additionally, the “Isolation Host” field does not always contain a bacterial host. In some cases, the animal or environmental source from which the phage was isolated is recorded, for example *Ciona robusta* (a sea squirt, n=257), *Hydrochoerus hydrochaeris* (capybara, n=148), *Homo sapiens* (n=147), and *Gopherus morafkai* (desert tortoise, n=119) all of which are clearly not hosts for bacteriophages. In other cases, the field is left blank. Clearer guidance for submitters on what this field should contain would improve the utility of this metadata.

### GenBank divisions blur the boundaries of bacterial and archaeal viruses

INPHARED originally used complete genomes from cultured isolates from GenBank, this focus continues with the inclusion of a small number of genomes from uncultured isolates. These are included due to their high level of interest within the phage genomics community, their representation of ICTV groupings, and extreme sizes. Examples include members of the *Crassvirales*^42–44^ and the Lak phages of the order *Grandevirales*^45–47^. Of the 28,777 genomes, 99.7% carry the PHG GenBank division. The remaining 71 carry the ENV division and one carries the VRL division. The “inclusion list” option allows the addition of cultured phage genomes that have inadvertently been submitted to environmental dataset (“ENV”) and not automatically identified.

The dataset also contains 209 genomes from archaeal hosts (e.g. *Sulfolobus*, *Halorubrum*, *Haloarcula*), all of which carry the PHG GenBank division. Whilst these are not bacteriophages, there is no dedicated GenBank division for archaeal viruses; some are designated PHG, others VRL, with no clear convention. Taxonomically, there is partial overlap between archaeal viruses and phages: 72 of the 209 archaeal virus genomes belong to the class *Caudoviricetes*, which is shared with the vast majority of cultured phages and four belong to *Tectiliviricetes* that is similarly shared between bacterial and archaeal viruses. The remainder belong to archaeal-specific classes including *Tokiviricetes* (23), *Huolimaviricetes* (20), *Laserviricetes* (8), and *Belvinaviricetes* (2), with 80 unclassified. Similarly, 12 archaeal-specific families are present in the dataset (e.g. *Fuselloviridae*, *Rudiviridae*, *Pleolipoviridae*, and *Bicaudaviridae*). In practice, archaeal viruses are frequently included alongside phages in viromics and comparative genomics analyses, but the lack of a standardised GenBank designation makes it difficult to distinguish them programmatically. A clearer system of GenBank divisions that indicates the host domain (e.g. VRL-Bac, VRL-Arc, VRL-Euk) could improve consistency for both submitters and downstream users. In the meantime, we have clearly identified these viruses within the INPHARED dataset.

### Host bias persists despite broadening diversity

The host bias identified in the 2021 INPHARED data persists, with the top 30 host genera accounting for 86.5% (2021) and 85.5% (2026) of genomes with an identified host, although their composition has shifted (Figure 2). The host bias can skew reference databases when developing bioinformatic tools, as if not accounted for the over presentation can lead to tools/models that are biased towards the dominant well characterised hosts.

**Figure 2.**
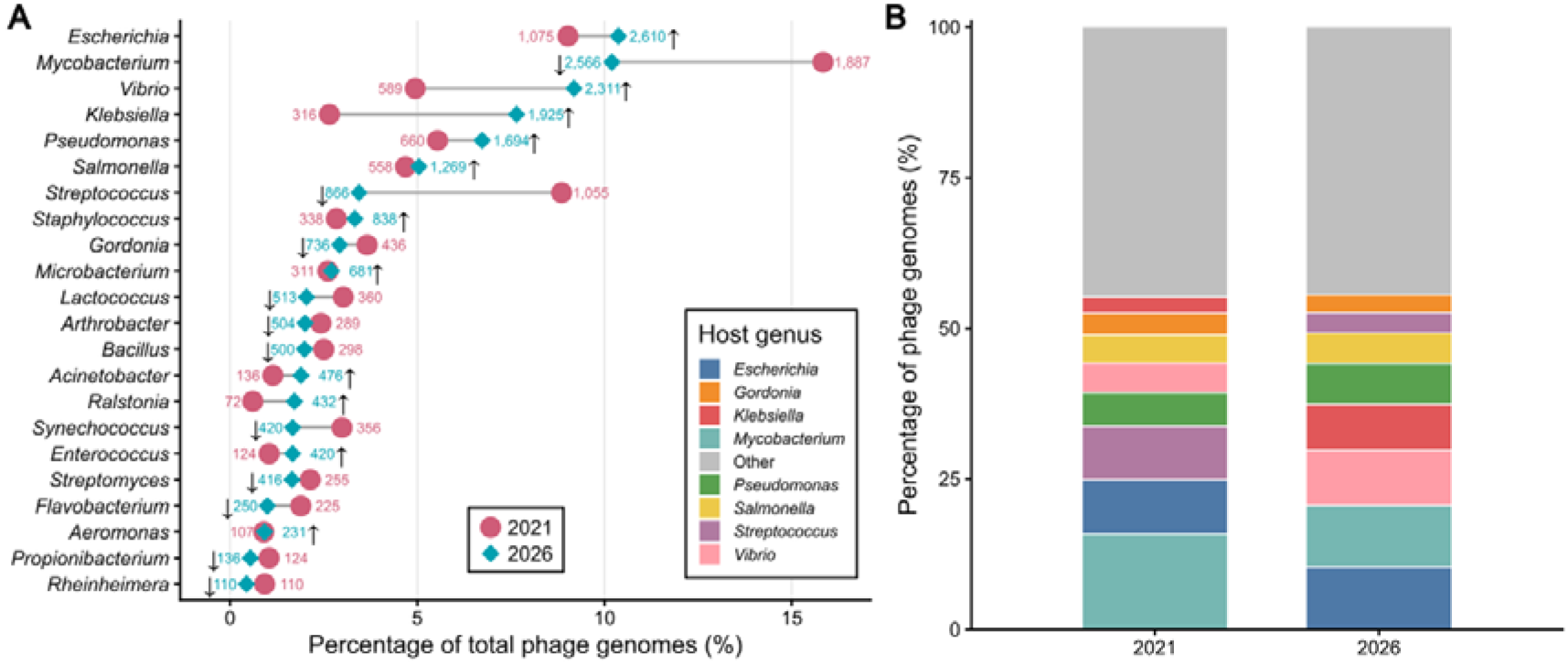
Shifts in most common isolation hosts. (**A**) Dumbbell plot showing percentage of total phage genomes isolated across the top 20 most common isolation hosts for 2021 and 2026 datasets (22 genera shown in total). Plotted values are percentage counts, but number of genomes is also displayed. Arrows indicate direction of proportional change. Y-axis is ordered by 2026 values. (**B**) Stacked barchart showing the proportional representation of the top 7 most common host genera for 2021 and 2026 datasets (8 genera shown in total), with all remaining genera grouped as “Other”. Bars independently ordered in descending order for their year.

*Escherichia* is now the most common host genus (n=2,610; +143%), displacing *Mycobacterium* (n=1,887; +36%), likely reflecting the broadening of the SEA-PHAGES programme to other genera within *Actinomycetota*^48^. The most pronounced increases were seen for *Klebsiella* (n*=*1,925; +509%) and *Vibrio* (n=2,311; +292%). Other notable increases included *Pseudomonas* (n=1,694; +157%), *Salmonella* (n*=*1,269; +127%), *Staphylococcus* (n*=*838; +148%), *Ralstonia* (n*=*432; +500%), *Acinetobacter* (n*=*476; +250%), and *Enterococcus* (n*=* 420; +239%).

The composition of newly sequenced genomes varies across these hosts. New *Escherichia* phages show the greatest genus-level diversity (96 genera), whilst *Salmonella* phages are the most taxonomically well-characterised of the fast-growing hosts, with only 34% unclassified at the family level. In contrast, new *Vibrio* phages are largely unclassified (91% at both family and genus level) and 72% lack a reported isolation host. Growth in *Klebsiella*, *Pseudomonas*, *Acinetobacter*, and *Staphylococcus* phages is broadly consistent with the expansion of phage therapy research, where these are priority pathogens^49^.

Enterobacteria has been eliminated as a host designation (487 in 2021), as it is not a valid genus. Of the 487 original accessions, 480 are present in the 2026 dataset: 474 reclassified to “Unspecified” and six to *Escherichia*. The remaining seven were removed through curation. *Streptococcus* showed an apparent decrease (1,055 to 866), but this is entirely attributable to the removal of 348 satellite phages described above; 159 new *Streptococcus* phage genomes were added to the 2026 dataset.

Host diversity has expanded, with 333 unique host genera now represented compared to 236 in 2021 (97 new genera). Notable additions include *Curtobacterium* (n=51), *Porphyromonas* (n=29), and *Phocaeicola* (n=25). Despite this growth, the overall shape of the host distribution remains largely unchanged, with the database dominated by a small number of clinically and agriculturally relevant genera.

However, the accuracy of host assignments is further complicated by changes to bacterial taxonomy itself. Several host genera present in the database have been partially or wholly reclassified in recent years, and this is inconsistently reflected in the metadata. *Lactobacillus*, which was split into 25 genera in 2020^50^, remains the host designation for 98 phages, compared to only 39 assigned to the new genera (e.g. *Lacticaseibacillus*, *Latilactobacillus*). In some cases, the “Isolation Host” field uses the updated genus name while the Host field in INPHARED retains the old name (which is taken from the phage name), creating inconsistencies within the same record. Similarly, all 172 *Clostridium* phages retain the old genus name, despite the reclassification of *C. difficile* to *Clostridioides difficile* in 2016^51^; a separate set of 21 phages are assigned to *Clostridioides*. *Mycobacterium* (2,566 genomes) likewise far outnumbers *Mycolicibacterium* (21), following the division of the genus in 2018^52^. The 166 phages hosted by *Shigella*, a genus that is genomically nested within *Escherichia* ^53^, present a further example of this taxonomic ambiguity.

### Genomic taxonomy is expanding but still has gaps

Between the 2021 and 2026 snapshots, the ICTV Bacterial Viruses Subcommittee introduced substantial changes to phage taxonomy. The morphology-based families *Myoviridae*, *Siphoviridae*, and *Podoviridae*, along with the order *Caudovirales*, were abolished (ratified 2022), after the families were shown to be to be polyphyletic^11^. In parallel, the broader adoption of a mega-taxonomic framework for virus classification introduced higher taxonomic ranks including class, phylum, kingdom and realm^12^.

The 2026 dataset now includes higher taxonomic ranks absent in 2021 as result of the introduction of this mega-taxonomy. There are 19 unique orders, 10 classes, nine phyla, and nine kingdoms represented. At the class level, *Caudoviricetes* dominates (78.3%), followed by *Malgrandaviricetes* (9.5%), with 9.9% unclassified. As a result of shifts in the taxonomic framework, the proportion of genomes assigned to a family dropped from 95.6% (2021) to 45.9% (2026), reflecting the removal of the three former morphological based families which previously dominated assignments (Figure 3). Although the number of unique families has increased from 28 to 120, many genomes have yet to be reassigned. This is most apparent for the former *Siphoviridae* family, where 86.1% of genomes previously assigned to this family remain unclassified at the family level in 2026, compared to 61.3% for *Podoviridae* and 43.2% for *Myoviridae*.

**Figure 3.**
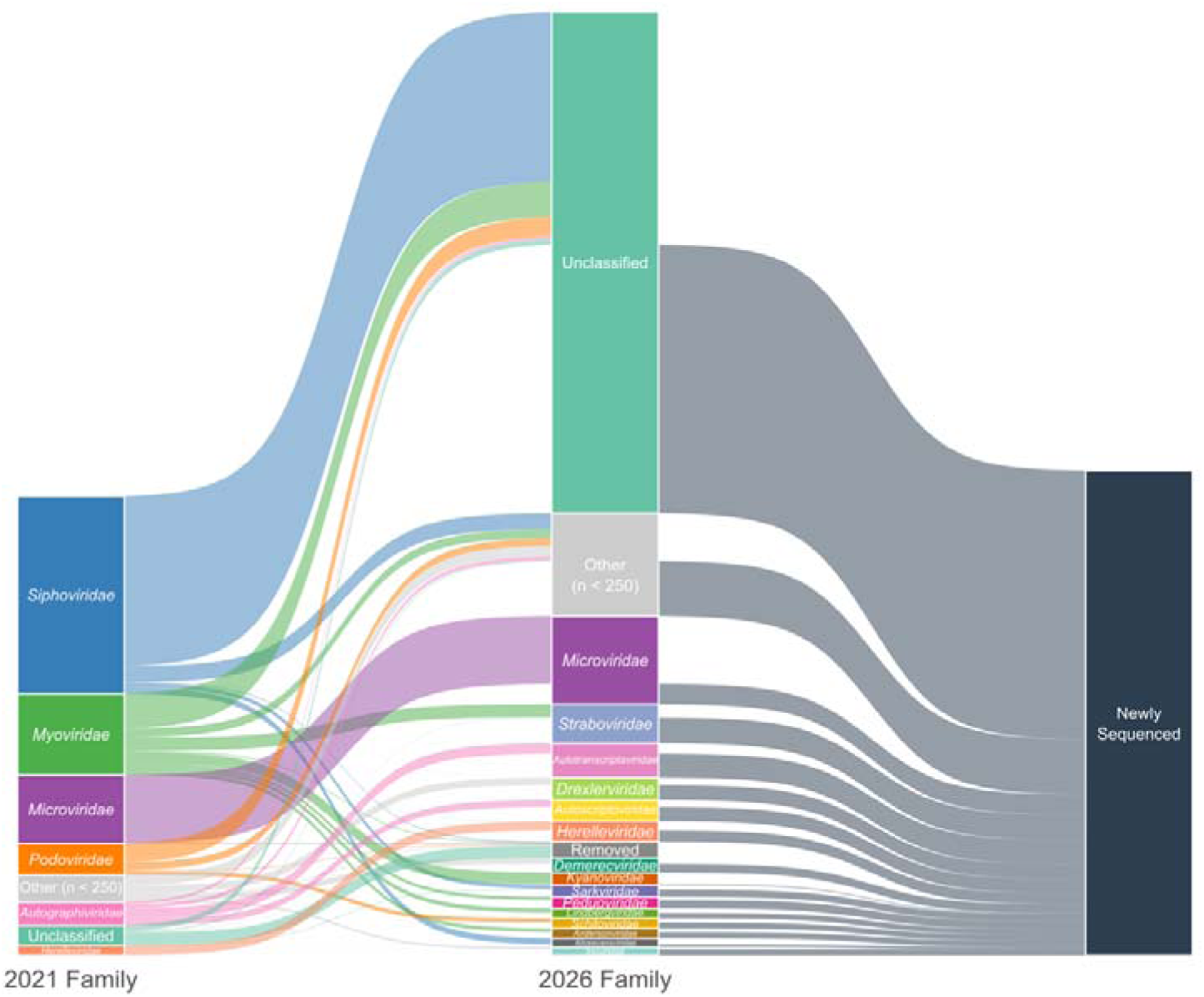
Overhaul of bacteriophage taxonomy. Alluvial plot showing Family level assignments in 2021 and 2026 with connections indicating how taxonomy has changed in these five years. Newly sequenced genomes in the interval period are also shown on the right of the plot. Families with < 250 members are collapsed into “Other”. Bars are scaled to number of genomes.

At the genus level, the number of unique genera has increased from 708 to 1,717. The proportion of genomes assigned to a genus has increased from 60.7% to 66.2%, suggesting that genus-level classification has kept better pace with the influx of new genomes than family-level classification. As the definitions of viral genera and species in the class *Caudoviricetes* are now clearly defined by nucleotide similarity, it allows for the rapid and automated identification of existing strains, species, and genera. As such, we have incorporated *taxMyPhage* results into the INPHARED database, allowing users to rapidly identify all phages within a genus or species, and identify novel species or genera within the dataset^30^. The inclusion of *taxMyPhage* results classified an additional 2,469 genomes at the genus level that were unclassified by the extraction of GenBank data alone, and revised genus-level assignments for 146 genomes where the two sources disagreed. These genus assignments also resolved higher-rank taxonomy, classifying an additional 1,409 genomes at the family level.

### Genomic properties remain stable

Despite the dataset doubling in size, the overall genomic properties of sequenced phages have remained broadly stable, suggesting consistent underlying characteristics.

Mean genome length increased modestly from 59.9 kb to 65.0 kb (median from 44.7 kb to 47.0 kb). A Wilcoxon rank-sum test confirmed this shift to be statistically significant (p < 10⁻¹D), but the effect size is small (rank-biserial r = −0.063; median difference = 2.29 kb). The shift is more pronounced when comparing the 2021 genomes to only those newly added in 2026 (median difference = 3.34 kb, r = −0.112), indicating that newly sequenced phages tend to be somewhat longer. Within individual realms, median lengths were essentially unchanged (Figure 4A), with *Duplodnaviria* showing a bimodal distribution at ∼42 kb and ∼165 kb that reflects widely sampled T7- and T4-like genomes, respectively. The size range has expanded slightly at extremes (3,143–642,428 bp in 2021 to 1,761–735,411 bp in 2026). So-called “jumbo-phages” (≥200 kb) remain proportionally stable at ∼2.2–2.3% of all genomes, doubling only in absolute number. GC content and coding capacity has remained essentially unchanged (means ∼ 48% GC and ∼90% coding capacity) and the proportion of genomes carrying at least one tRNA has increased slightly from 32.2% to 34.0%. CDS counts increased from 86.7 to 94.3 (median 66 to 69) consistent with the modest increase in genome length.

**Figure 4.**
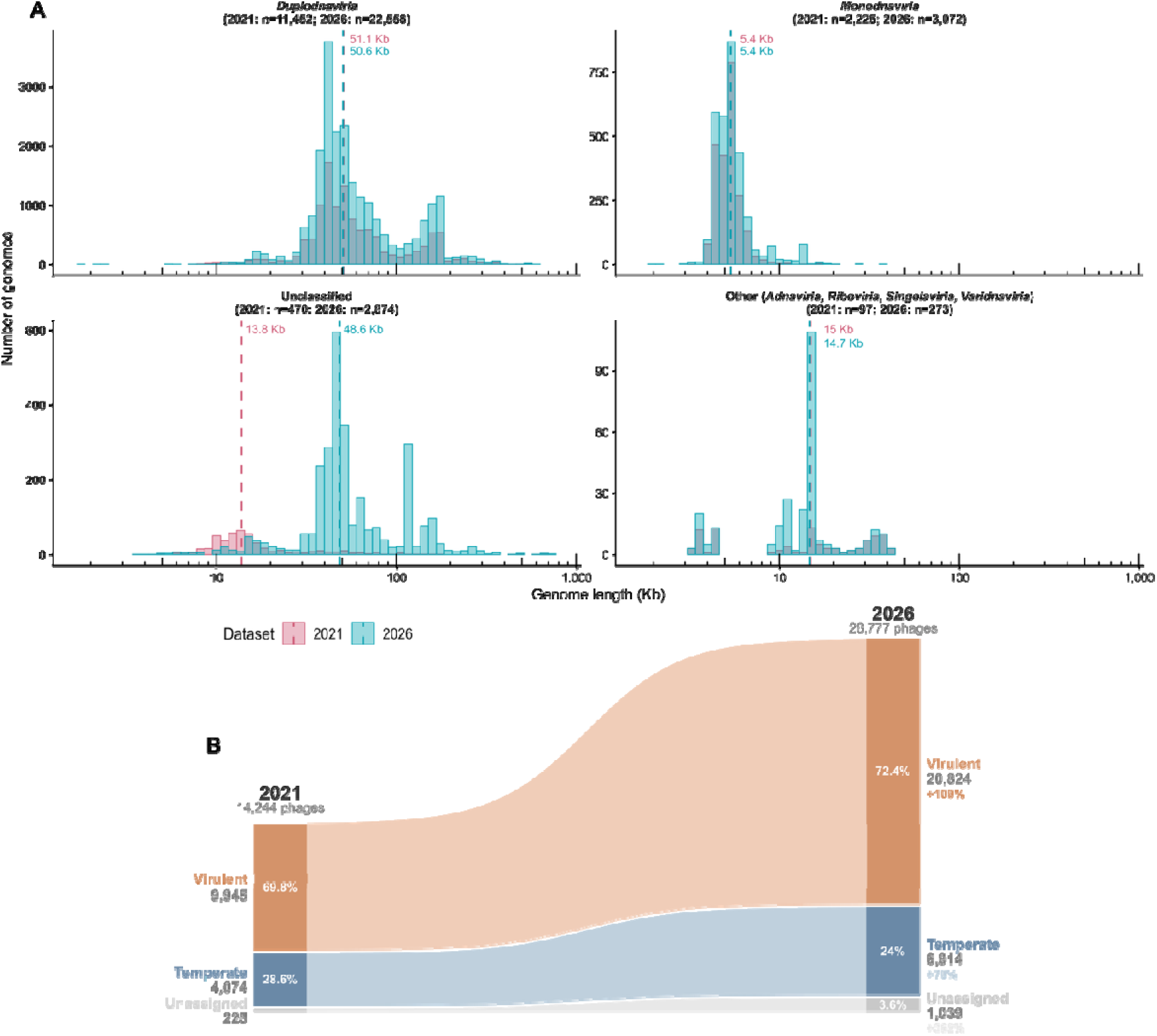
Genomic Features of Phages. (**A**) Histograms comparing genome lengths in the 2021 and 2026 datasets, faceted by Realm (with *Adnaviria*, *Riboviria*, *Singelaviria*, and *Varidnaviria* combined for simplicity of plotting). Dashed lines and annotation indicate the median value. (**B**) Comparison of predicted lifestyles of phages across the 2021 and 2026 datasets.

Virulent (or “obligately lytic”) phages, predicted using PhaTYP^31^, remain dominant in our database and have increased lightly (69.8% to 72.4%), while temperate phages decreased proportionally (28.6% to 24.0%) despite increasing in absolute numbers by 70% (Figure 4B). Notably, the unassigned category grew substantially (225 to 1,039; ∼362% increase) suggesting an increasing fraction of newly sequenced cannot be confidently classified by lifestyle prediction tools used in INPHARED. GenBank does not currently include a dedicated field for phage lifestyle; addition of such a field would allow submitters to record experimentally determined lifestyle, where known, and reduce reliance on prediction tools.

### Anti-defence systems are widespread and linked to genome size

With increasing interest in phage therapy and the selection of suitable phages, we have incorporated the automated identification of both defence and anti-defence systems in phage genomes using DefenseFinder^17^. Analysis of the current dataset identified 6,454 genomes (22.4%) carrying at least one anti-defence system, encompassing 16 system types and 97 subtypes (Fig 5). Anti-RM was the most commonly detected type (2,447 genomes), followed by Anti-CBASS (2,019) and Anti-TA (1,742). Anti-CRISPR systems, despite being the most widely studied class of anti-defence, ranked sixth (558 genomes), behind NADP (1,405) and Anti-Thoeris (760).

**Figure 5.**
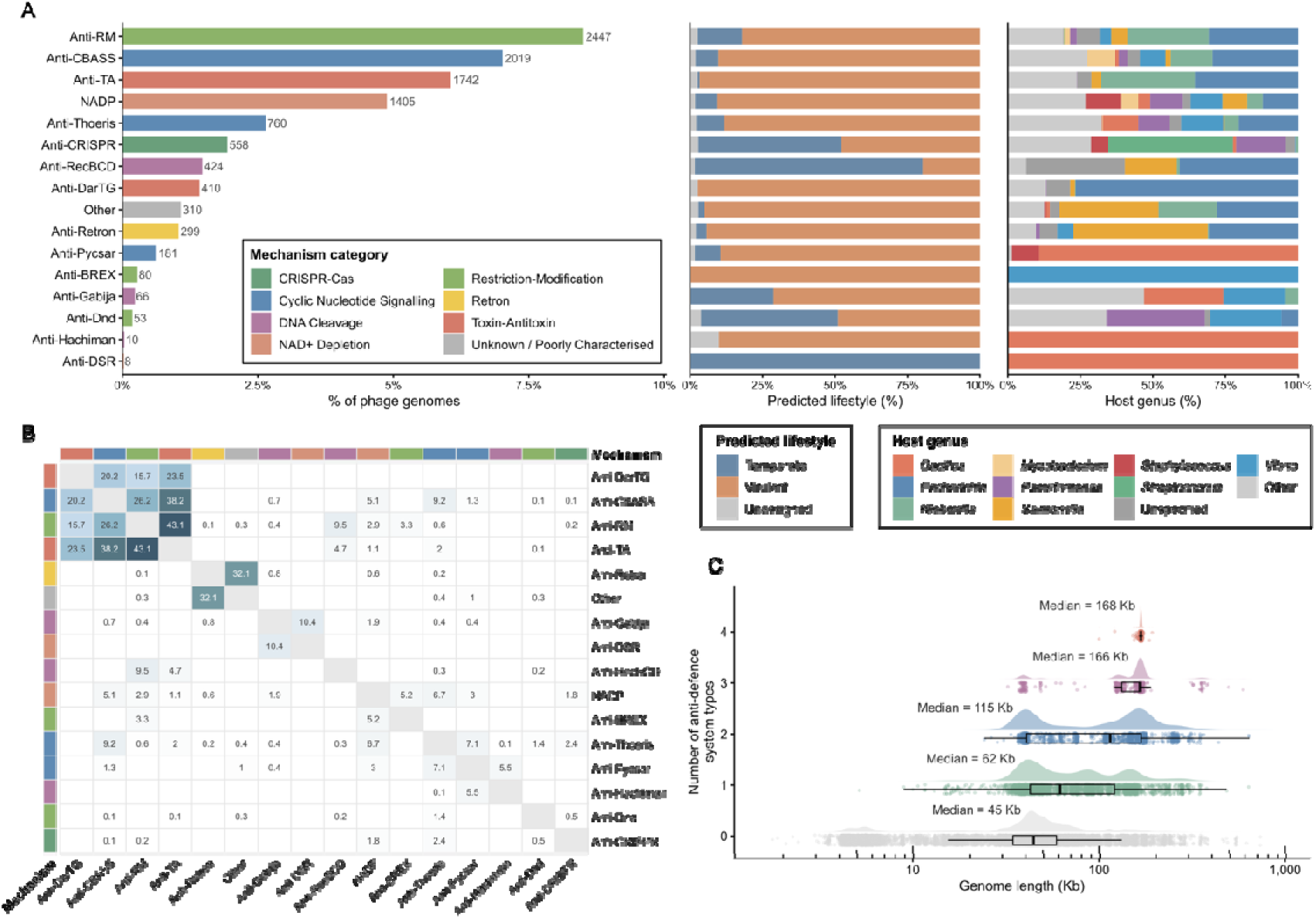
Anti-defence system carriage. (**A**) Barplot showing anti-defence systems encoded on phage genomes, colour coded by mechanism category (left), with predicted lifestyle of carrier phages (middle) and host genus (right) for each anti-defence system. Only the top ten most frequent host genera in the plot are shown (including “Unspecified”), with all others being collapsed into “Other”. (**B**) Co-occurrence heatmap of anti-defence systems with displayed value indicating percentage probability of two systems occurring together. (**C**) Raincloud plot of genome lengths for phages with 0, 1, 2, 3 or 4 anti-defence system types encoded on their genome.

Anti-defence carriage was strongly associated with genome size (Figure 5). Genomes carrying at least one anti-defence system had a median length of 88,858 bp compared to 44,651 bp for those without (Wilcoxon rank-sum test, p < 10⁻¹D). The number of anti-defence systems per genome was positively correlated with genome length (Spearman’s rho = 0.39, p < 10⁻ ¹D). Most carrier genomes encoded a single system (3,404/6,454; 52.7%), with 1,516 encoding two and 750 encoding three. However, 395 genomes (6.1%) carried four distinct anti-defence types simultaneously, and 2,857 genomes (44.3% of all carriers) encoded at least two distinct types.

Anti-defence carriage varied substantially across phage families. Members of the family *Straboviridae* had the highest carriage rate among well-represented families, with 1,206 of 1,212 genomes (99.5%) encoding at least one anti-defence system. Other families with high carriage rates included *Rudiviridae* (23/23; 100%; Archaeal viruses), *Andersonviridae* (289/305; 94.8%), *Vilmaviridae* (98/104; 94.2%), *Jeanschmidtviridae* (33/39; 84.6%), and *Autotranscriptaviridae* (835/1,070; 78.0%). At the other end of the spectrum, *Microviridae* showed near-complete absence (1/2,736; 0.04%), consistent with the small genome sizes of this family. *Peduoviridae* (0/357), *Tectiviridae* (0/101), *Salasmaviridae* (0/61), and *Fiersviridae* (0/39) also lacked detectable anti-defence systems entirely.

Several families showed highly conserved anti-defence profiles. All 289 anti-defence systems detected in *Andersonviridae* were NADP, all 163 in *Aliceevansviridae* were Anti-CRISPR, and 99 of 100 in *Autosignataviridae* were Anti-RM. Other families displayed more diverse profiles: *Herelleviridae* showed a broad distribution across NADP (44.0%), Anti-Pycsar (25.4%), Anti-Thoeris (17.9%), and Anti-CRISPR (6.2%), and *Demerecviridae* was dominated by Anti-Retron (50.4%) and the orf148 anti-defence protein (37.4%).

Within *Straboviridae*, the distribution of systems per genome was bimodal, with peaks at approximately three and six to seven systems per genome. Based on these peaks, genomes were divided into a low group (1–4 systems; n = 637) and a high group (5–10 systems; n = 569). The two groups did not differ substantially in genome length (median 170,286 bp and 167,921 bp) but mapped clearly onto genus-level taxonomy. The high group was almost exclusively represented by the subfamily *Tevenvirinae* (568/569), dominated by the genera *Tequatrovirus* (405/569) and *Mosigvirus* (129/569), and predominantly isolated on *Escherichia* (427/569). These genomes carried a characteristic gene complement including *acb1, acb2, dmd, tifa, arn, adfa, ipi, ipii,* and *dam*. Anti-DarTG (*adfa*) was present in 391/569 high-group genomes but entirely absent from the low group, effectively serving as a marker of this *Tevenvirinae* lineage. The 13 most heavily armed genomes in the entire dataset, each carrying 10 distinct anti-defence systems, were all *Tevenvirinae* members with genome lengths of 166–170 kb.

The low group was taxonomically more diverse, spanning three subfamilies (*Tevenvirinae* with 267/637, *Twarogvirinae* with 48/637, and *Emmerichvirinae* with 12/637) and 310 genomes unclassified at the subfamily level. The *Tevenvirinae* genomes in the low group comprised largely different genera to those in the high group, dominated by *Jiaodavirus* (137/267), *Dhakavirus* (30/267), and *Karamvirus* (30/267), whilst *Tequatrovirus* and *Mosigvirus* contributed only 10 and 1 respectively. These phages infected a broader range of hosts, with *Klebsiella* (250/637), *Aeromonas* (51/637), *Acinetobacter* (49/637), and *Enterobacter* (48/637) among the most common. Their anti-defence complement was distinct: *acb1, alt,* and *dam* were the dominant subtypes, and Anti-Thoeris and NADP were present in this group but absent from the high group. It should be noted that many of the anti-defence systems enriched in the high group (e.g. *dmd, alt, arn, ipi, and ipii*) were originally characterised in coliphage T4^16^, the archetypal *Tequatrovirus*. The magnitude of the difference between the two groups may therefore be partially inflated by a detection bias in DefenseFinder’s models, which are necessarily built from previously characterised systems.

Co-occurrence analysis across all genomes revealed several strong associations between anti-defence types. Anti-BREX co-occurred with Anti-RM in all 80 genomes carrying it, and with NADP in 91.2%; this is consistent with the complementary roles of BREX and RM in bacterial defence, which together protect against both modified and non-modified invading DNA. The three most abundant types (Anti-CBASS, Anti-RM, and Anti-TA) frequently co-occurred, with 72.4% of Anti-TA-carrying genomes also encoding Anti-RM and 59.6% also encoding Anti-CBASS, largely reflecting the *Straboviridae* contribution. These co-occurrence patterns likely reflect the defence system repertoires of the hosts these phages infect; bacterial defence systems are known to co-occur non-randomly and act synergistically^54^, and phages infecting a given host would require counters to its specific combination of defences.

When normalised by host genus, the proportion of phages carrying anti-defence systems was highest for *Shigella* (120/166; 72.3%), *Brevibacillus* (7/11; 63.6%), *Cronobacter* (33/55; 60.0%), *Yersinia* (112/192; 58.3%), *Citrobacter* (39/67; 58.2%), and *Sulfolobus* (43/77; 55.8%; Archaeon). Among the most well-sampled hosts, *Klebsiella* (1,014/1,925; 52.7%), *Escherichia* (1,283/2,610; 49.2%), *Erwinia* (101/208; 48.6%), *Aeromonas* (109/231; 47.2%), and *Bacillus* (228/500; 45.6%) all had substantial carriage rates. In contrast, *Mycobacterium* phages had a notably low rate (265/2,566; 10.3%) despite being the second most sampled host, likely reflecting both their smaller genome sizes and the predominance of temperate phages in this group. Similarly, *Gordonia* (36/736; 4.9%), *Lactococcus* (7/513; 1.4%), and *Streptomyces* (1/416; 0.2%) showed very low carriage.

### Defence system carriage is rare and enriched in temperate phages

The incorporation of DefenseFinder results identified bacterial defence systems within phage genomes, with 785 genomes (2.7%) encoding at least one system, an order of magnitude lower than anti-defence carriage (Figure 6)^17^. Defence systems encoded within phage genomes are thought to function primarily in superinfection exclusion, protecting the lysogen from subsequent infection by competing phages. Consistent with this, defence systems were enriched in temperate phages (599/6,914; 8.7%) compared to virulent phages (172/20,824; 0.8%), whereas anti-defence carriage showed the opposite pattern (16.4% vs 24.9%). Despite their lower prevalence, 96 defence system types and 112 subtypes were detected, with most genomes encoding a single defence system (668/785; 85.1%). The most common defence system was restriction-modification (n=184) followed by AbiAlpha (n=64), AbiD (n=54), CRISPR-Cas (n=49; predominantly Class 1 Subtype I-F,n= 46), VP1839 (n=35), PD-Lambda-1 (n=31), Retron (n=29), FS_Sma (n=26), and MMB_gp29_gp30 (n=23). Many of the recently described defence systems were also represented, including Septu (17), Kiwa (10), Gabija (3), and Pycsar (3). In contrast to anti-defence systems, defence system carriage was not associated with larger genomes. Most detected defence systems were complete (83.8%). Among incomplete systems, the CRISPR-Cas system was most represented, as there are no instances where it is found in full (n=242), while Restriction Modification system completeness varied widely (0.17 to 1; n=360; Table 1). The variability is likely driven by the required components, from simple Type IV systems (1 gene) to Type I systems (3 genes)^55^.

**Figure 6.**
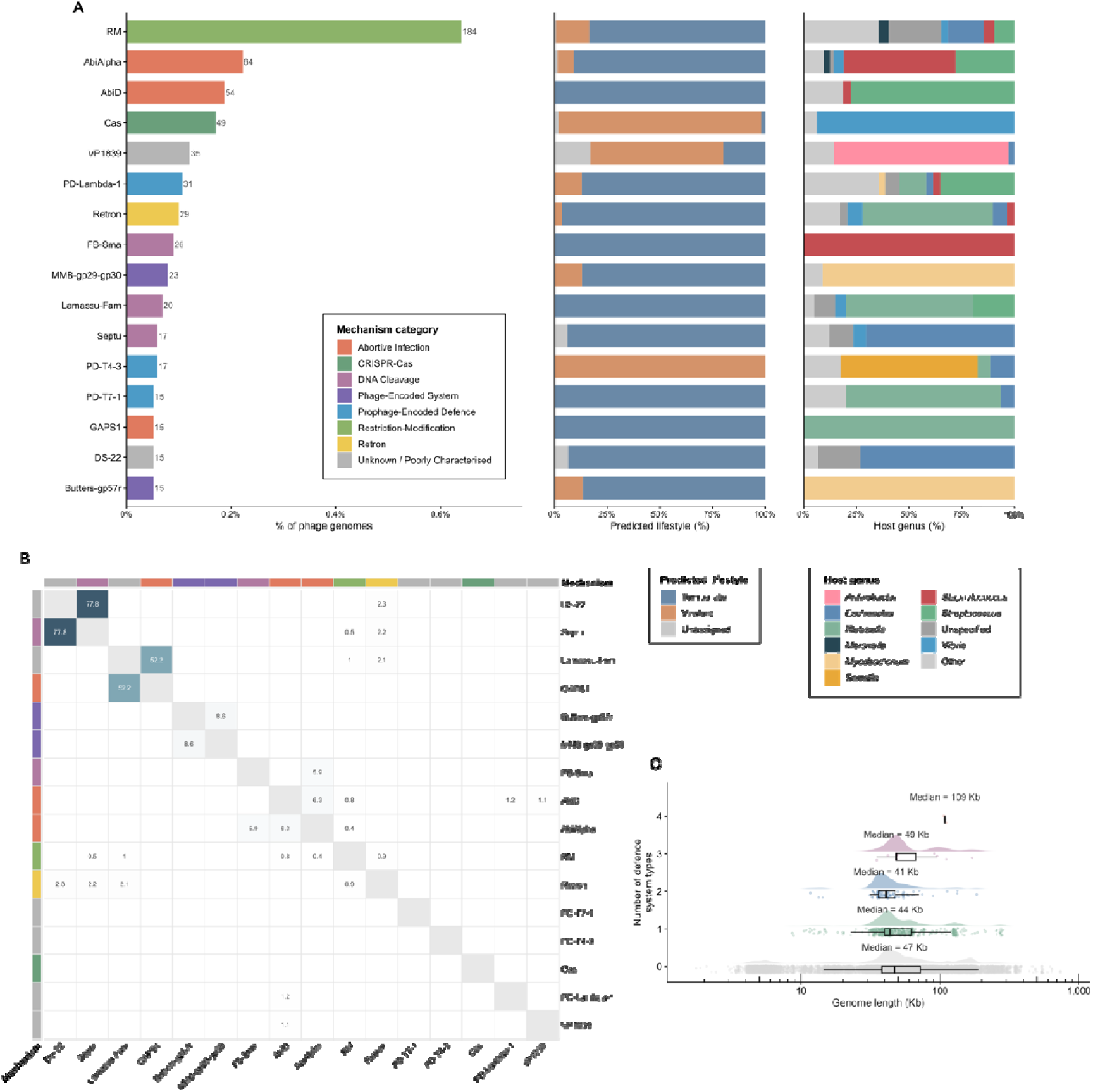
Defence-system carriage. (**A**) Barplot showing defence systems encoded on phage genomes, colour coded by mechanism category (left), with predicted lifestyle of carrier phages (middle) and host genus (right) for each defence system. Only systems for which ≥ 15 genomes are predicted to carry are shown. Only the top ten most frequent host genera in the plot are shown (including “Unspecified”), with all others being collapsed into “Other”. (**B**) Co-occurrence heatmap of defence systems with displayed value indicating percentage probability of two systems occurring together. (**C**) Raincloud plot of genome lengths for phages with 0, 1, 2, 3 or 4 defence system types encoded on their genome.

**Table 1:** Completeness of defence systems found in bacteriophage genomes – 11 system types with at least one partial detection, ordered by ascending order of completeness. Calculated from the sys_wholeness parameter representing a faction of expected system components detected by the DefenseFinder^17^, where 1.00 represents a fully complete system.

| *System* | *Total*<br>*instances* | *Complete*<br>*(n)* | *Complete*<br>*(%)* | *Mean*<br>*wholeness* | *Min* | *Max* |
| --- | --- | --- | --- | --- | --- | --- |
| *Cas* | 242 | 0 | 0 | 0.825 | 0.25 | 0.833 |
| *DRT* | 1 | 0 | 0 | 0.25 | 0.25 | 0.25 |
| *DS-4* | 14 | 0 | 0 | 0.4 | 0.4 | 0.4 |
| *FS_HsdR_like* | 2 | 0 | 0 | 0.667 | 0.667 | 0.667 |
| *PrrC* | 12 | 0 | 0 | 0.625 | 0.5 | 0.75 |
| *SDIC3* | 7 | 0 | 0 | 0.714 | 0.6 | 0.8 |
| *Toutatis* | 3 | 0 | 0 | 0.5 | 0.5 | 0.5 |
| *Sucellos* | 6 | 2 | 33.3 | 0.667 | 0.5 | 1 |
| *Lamassu-Fam* | 47 | 21 | 44.7 | 0.816 | 0.667 | 1 |
| *RM* | 360 | 286 | 79.4 | 0.856 | 0.167 | 1 |
| *Retron* | 59 | 57 | 96.6 | 0.983 | 0.5 | 1 |

Abortive infection (ABi) systems were the most prevalent mechanism in ∼26% of defence carrying genomes. Given Abi typically results in host cell death and limits phage propagation^56^, this raises questions on how phage-encoded ABI are regulated and avoid self-inhibition. Among the 117 phages carrying more than one defence system, 64 unique pairwise combinations were observed. The top pairs were DS-22 + Septu (n=14), GAPS1 + Lamassu-Fam (n=12), and DS-16 + PD-T7-1 (n=11). The three *Lambdavirus*-associated pairs, DS-22 + Septu, RexAB + Septu, and DS-22 + RexAB, represent the clearest evidence of a conserved multi-system module, with many *Lambdavirus* phages carrying all three simultaneously. Among the 12 systems appearing in five or more multi-system phages, two distinct clusters emerged that appear mutually exclusive: an Abi cluster (AbiAlpha, AbiD, FS_Sma) and a prophage/DNA-cleavage cluster (DS-16, DS-22, GAPS1, Lamassu-Fam, PD-T7-1, RexAB, Septu). No member of either cluster ever appears alongside a member of the other, suggesting these represent distinct ecological or taxonomic niches rather than randomly assembled arsenals. RM sits outside both clusters, pairing with RloC and Retron but never with any DS-series or Abi systems.

Only a small number of classified bacteriophage families were represented amongst defence-carrying phages, namely *Peduoviridae*, *Chimalliviridae*, and *Duneviridae*. *Peduoviridae* has the most diversity, with 24 distinct system types across 79 phages. In contrast the *Chimalliviridae* family carry only one type of system (PD-T4-3, n=12). At genus level, six genera with ≥5 phages carried just one defence system type: *Oslovirus* (RM, n=49), *Mohonavirus* (Cas, n=43), *Bridgettevirus* (VP1839, n=7), *Nanditavirus* and *Richievirus* (both VP1839, n=5), and *Spinunavirus* (AbiD, n=5). These are strikingly uniform, with every phage in the group carrying the same defence system. *Lambdavirus* carries the most per phage on average at 2.5 systems/phage, with most phages carrying all three of DS-22, Septu, and RexAB simultaneously. Most classified families carried no detectable defence systems, including *Straboviridae* (0/1,212), *Autotranscriptaviridae* (0/1,070), and *Drexlerviridae* (0/692). When normalised by host genus, *Streptococcus* phages showed the highest defence carriage rate among well-sampled hosts (111/866; 12.8%), followed by *Staphylococcus* (78/838; 9.3%), *Burkholderia* (14/197; 7.1%), *Arthrobacter* (29/504; 5.8%), and *Clostridium* (9/172; 5.2%). *Streptococcus* and *Staphylococcus* are both genera in which temperate phages carrying accessory genes are well documented^57,58^, and the enrichment of defence systems in their phages is consistent with a role in superinfection exclusion. In contrast, several well-sampled hosts had no defence-carrying phages at all, including *Aeromonas* (0/231), *Erwinia* (0/208), *Xanthomonas* (0/199), and *Streptomyces* (0/416).

A total of 163 genomes (0.6%) carried both defence and anti-defence systems, predominantly among temperate phages (99/163; 60.7%). This dual carriage was dominated by *Vibrio* phage ICP1 (40 genomes), which encodes its own CRISPR-Cas system to counter *V. cholerae* phage-inducible chromosomal island-like elements (PLEs), alongside anti-defence systems^59,60^. Lambda-like *Escherichia* phages also featured prominently, consistent with the RexAB superinfection exclusion system first described in Lamda^61^. Several *Streptococcus* phages, phage-plasmids P1 and P7, and a small number of *Chimalliviridae* members also carried both system types. The coexistence of both defence and anti-defence systems within the same genome likely reflects the dual pressures on temperate phages: overcoming host defences during infection while protecting the lysogen from competing phages after integration.

### ARG and virulence factor carriage remains low

In the 2026 INPHARED dataset antimicrobial resistance genes (ARGs) and virulence factors (VFs) have increased in absolute numbers (ARGs: 43 to 70; VFs: 235 to 567) but decreased slightly in proportion to dataset growth (ARGs: 0.30% to 0.24%; VFs: 2.42% to 1.97%). The diversity of genes has expanded from 32 to 43 for ARGs and from 50 to 140 for virulence factors, with all genes present in 2021 retained in 2026.

The most common ARGs remain macrolide resistance genes *mef*(A) and *msr*(D) (21 and 17 genomes, up from 14 and 12 respectively). *Streptococcus* remains the dominant host, though new hosts include *Acinetobacter*, *Enterococcus*, and *Pseudomonas.* Notably Escherichia phage P1 (accession MH445380) was found to carry eight ARGs spanning seven resistance classes, highlighting its potential role in ARG dissemination^62^.

For virulence factors (VFs), the most notable change was a substantial increase in Shiga toxin-encoding phages. *stx2A* increased from 73 to 185 genomes and stx2B from 72 to 184, with the majority of new detections in phages with an unspecified host. As with the broader dataset, the 18 phages previously assigned to “Enterobacteria” have been reclassified to “Unspecified”, contributing to the 201 VF-carrying phages now lacking a genus-level host designation. *Staphylococcus* VF-carrying phages have nearly doubled (57 to 101). Five new host genera for VF-carrying phages appear: *Listeria* (5 genomes; internalins and immune modulators), *Enterococcus* (4 genomes; capsule and cytolysin operons), *Campylobacter*, *Helicobacter*, and *Salmonella*. As noted for the *Enterococcus* and *Listeria* phages discussed above, several of these entries involve multiple accessions sharing the same phage name with markedly different genome sizes, which may reflect fragmented assemblies deposited as separate entries or mis-assigned bacterial assemblies deposited to GenBank as phage genomes.

The taxonomic changes described above are starkly reflected in these data. In 2021, ARG-carrying genomes were assigned to *Siphoviridae*, *Myoviridae*, and *Podoviridae*; in 2026, all 70 are unclassified at the family level. Similarly, 553 of 567 VF-carrying genomes are now unclassified at the family level, with only *Inoviridae* (n=110), *Peduoviridae* (n=2), and *Aliceevansviridae* (n=1) retaining assignments.

### Novel phages are underrepresented in quality reference databases

CheckV was used to assess genome quality and completeness in both datasets^32^ and is now included in the metadata of the INPHARED dataset. Most genomes were classified as Complete or High-quality, though this proportion decreased slightly from 94.5% (13,435/14,222) in 2021 to 92.3% 26,555/28,777) in 2026. The proportion of Complete genomes increased (10.8% to 14.5%) largely due to detection of terminal repeats. However, as transposon-based library preparation methods do not capture the physical termini of genomes, most complete phage genomes will lack detectable DTRs and will therefore not receive the CheckV Complete classification. Conversely, short-read assemblies from SPAdes can produce artefactual DTRs, and CheckV itself notes that terminal repeats can represent assembly artefacts.

The representation of INPHARED genomes within the CheckV reference database has declined as phage diversity has increased. In 2021, 92.3% of genomes shared ≥90% AAI with their closest CheckV reference, compared to 81.9% in 2026. Similarly, the proportion of genomes with less than 50% alignment fraction to their closest reference increased from 2.5% to 7.0%. The number of genomes receiving low-confidence completeness estimates increased from 122 to 894, with these genomes having a mean AAI of 44.2% and a mean alignment fraction of 21.7% to their closest reference. This suggests a growing proportion of sequenced phages are poorly represented in the CheckV database, and that lower completeness estimates or quality classifications for these genomes may reflect limited database coverage rather than genuinely incomplete sequences. Thus, we have included CheckV outputs as part of our manual curation process and allow users to determine for themselves the usefulness of a phage genome.

### Conclusions

Over the last five years, the number of complete phage genomes from cultured isolates has doubled, yet this expansion has not translated into a proportional increase in novel diversity. The redundant sequencing of well-studied phages continues to outpace new discovery, reinforcing persistent host bias and highlighting a continued reliance on the same narrow set of bacterial hosts. Addressing this imbalance remains a central challenge to the field, requiring targeted expansion of hosts phages are isolated against.

The transition to genome-based taxonomic framework represents a transformative and necessary shift but still remains a work in progress. The abolition of the morphology-based families has exposed substantial gaps in higher level classification, with many genomes unclassified at the family level. In contrast, genus-level classification has proven more scalable, driven by clearly defined genomic criteria and use of automated tools (e.g. *taxMyPhage*). Continued refinement of higher taxonomic ranks will be essential as the dataset of phage genomes continues to grow.

The widespread distribution of anti-defence systems, contrasted with the rarity and lifestyle bias of defence systems, highlights the complexity of phage–host interactions and the selective pressures shaping phage genomes. Such features are increasingly relevant for translation into phage therapy, where genomic data is becoming increasingly used to inform to predict the safety and host range of phages.

Several systematic issues continue to limit the utility of public datasets. Inconsistent metadata, particularly for host information, the absence of standardised fields for empirically derived lifestyle, and lack of distinct GenBank division for archaeal viruses all hinder large scale data analyses that will become increasingly pronounced as the number of genomes continues to grow. As such the need for well-maintained, standardised reference databases becomes ever more critical. INPHARED will continue to be updated and expanded, and we encourage the phage community to contribute to its ongoing curation.

## Supplementary Material

Supplementary Tables S1–S10 are available on Figshare at (https://doi.org/10.6084/m9.figshare.32043228).

- **Table S1.** Per-genome metadata for the 2021 INPHARED dataset (accession, description, taxonomic classification, genome properties, host etc.).
- **Table S2.** Per-genome metadata for the 2026 INPHARED dataset (as S1, with additional fields for the full taxonomic ranks and GenBank division).
- **Table S3.** PhaTYP lifestyle predictions for the 2021 dataset.
- **Table S4.** PhaTYP lifestyle predictions for the 2026 dataset.
- **Table S5.** DefenseFinder output for anti-defence systems encoded on phage genomes (2026 dataset only).
- **Table S6.** DefenseFinder output for defence systems encoded on phage genomes (2026 dataset only).
- **Table S7.** Antimicrobial resistance gene hits from Abricate (NCBI database) for the 2021 and 2026 datasets.
- **Table S8.** Virulence factor hits from Abricate (VFDB) for the 2021 and 2026 datasets.
- **Table S9.** CheckV genome quality assessment for the 2021 dataset.
- **Table S10.** CheckV genome quality assessment for the 2026 dataset.

